# Coupling a developmental promoter to CRISPR interference for Wnt pathway regulation in human pluripotent stem cells

**DOI:** 10.64898/2026.09.25.754435

**Authors:** Gyuhyung Jin, Sean P. Palecek

## Abstract

While directed differentiation of human pluripotent stem cells commonly relies on the timed delivery of extracellular factors, these uniform treatments often yield heterogeneous responses across cell populations. Linking intracellular gene regulation directly to an emerging developmental state offers a complementary strategy to coordinate these differentiation signals from within the cell. Here, we explored this approach by coupling a T/Brachyury promoter to CRISPR interference targeting CTNNB1, which encodes the canonical Wnt signaling mediator β-catenin. A T-promoter EGFP reporter line exhibited transiently increased activity during early differentiation, supporting the use of this promoter as a developmentally responsive input. We then combined the promoter with dCas9-KRAB and a CTNNB1-targeting guide RNA. Cas9-mediated integration was accompanied by indels at the CTNNB1 target site, whereas a Cas12a-mediated integration strategy yielded clones with no indels detected by ICE analysis. During differentiation, the selected T-dCas9-CTNNB1 clone showed reduced CTNNB1 expression and attenuated induction of Wnt-associated genes. Together, these findings provide a proof of concept for combining a developmental promoter with a programmable intracellular regulator and identify a strategy for separating circuit integration from unintended target-site editing. This modular approach provides a foundation for developing genetic interventions whose expression is linked to developmental state.

## 1. Introduction

Controlled differentiation of human pluripotent stem cells (hPSCs) is essential for producing defined cell populations for disease modeling, drug discovery, and regenerative medicine. Many differentiation protocols translate developmental principles into sequential exposure to growth factors, cytokines, and small molecules. By activating or inhibiting signaling pathways at successive lineage decisions, these extracellular interventions guide hPSCs toward desired fates [1,2]. Their effectiveness depends on both the administered stimulus and the state in which cells receive it. Thus, a central challenge is to coordinate a differentiation signal with the differentiation potential of the responding cell.

A uniform culture treatment does not necessarily produce a uniform cellular response. For example, in hPSC cultures, colony geometry and endogenous signaling can generate spatially distinct differentiation patterns, while cell density changes the paracrine environment and alters cellular responses to differentiation cues [3,4]. Furthermore, scaling differentiation introduces additional process variables: aggregate size, agitation, small-molecule concentration, and induction timing all influence differentiation efficiency and lineage specification in suspension culture [5]. These observations suggest that variation in signal exposure and interpretation can contribute to heterogeneous differentiation. They also motivate strategies that supplement extracellular control with regulatory mechanisms operating inside each cell.

Embryonic development provides a compelling model for such cell-autonomous regulation, as cells reliably coordinate cell-fate decisions despite variable microenvironments. In these developing systems, spatially and temporally changing signals are interpreted through intrinsic gene regulatory networks that couple transcription factors, feedback mechanisms, and cellular identity. In the vertebrate neural tube, for example, a transcriptional network converts dynamic Sonic Hedgehog signaling into distinct expression states, buffering against extracellular fluctuations [6]. While reconstructing an entire developmental network remains a formidable engineering challenge, recent synthetic biology approaches have shown that selected regulatory interactions can be successfully engineered within hPSCs. Initially, forced expression of lineage-defining transcription factors demonstrated that ectopic regulators could drive rapid differentiation [7], a strategy later refined by using CRISPR activation to turn on endogenous regulatory loci [8]. CRISPR interference (CRISPRi) extends this programmable toolkit toward targeted gene repression through catalytically inactive Cas9 fused to transcriptional repressor domains such as KRAB [9,10]. Furthermore, epigenome editors such as CRISPRoff provide persistent, division-tolerant transcriptional memory [11]. Together, these complementary tools provide a practical foundation for engineering individual regulatory connections within intracellular developmental programs.

While these synthetic tools provide several ways to alter intracellular gene regulation, linking their activity to a cell’s changing developmental state remains a separate design challenge. Synthetic lineage-control networks have coordinated transcription factor expression using an externally supplied chemical input, and inducible CRISPRi systems in human iPSCs use doxycycline to initiate repression [12,13]. Light-responsive CRISPR systems likewise allow externally scheduled transcriptional control [14]. Other strategies, including conventional constitutive short-hairpin RNA expression cassettes, provide persistent suppression without an intrinsic developmental timing mechanism [15]. Importantly, this limitation stems from expression design rather than RNA-mediated regulation itself. For instance, microRNAs can be transcribed by RNA polymerase II and are therefore compatible with regulated promoter inputs [16]. Together, these examples distinguish the choice of regulatory effector from the control of its expression. Building on this distinction, a developmental promoter could link effector expression to a transcriptional state that emerges during differentiation.

We therefore investigated whether a mesendoderm-associated T/Brachyury promoter could serve as an input for intracellular pathway regulation in hPSCs. Brachyury is responsive to Wnt/β-catenin signaling, and a murine T-promoter fragment has been used to report early differentiation in human embryonic stem cells [17,18]. We selected β-catenin, encoded by CTNNB1, as the regulatory target because canonical Wnt signaling serves as a key regulator of hPSC fate decisions, where its activation strongly induces T/Brachyury expression during early mesendodermal differentiation [1,2,17]. Coupling this Wnt-responsive promoter back to a CTNNB1-directed repressor creates an autonomous negative-feedback loop, providing a defined model to test whether intracellular regulation can modulate the very pathway that triggered it.

To evaluate this concept, we implemented a stepwise approach spanning circuit validation, assembly, and transcriptional characterization. Specifically, we first monitored the activity profile of the integrated T promoter using an EGFP reporter, established an integration workflow that limits unintended cleavage at the regulatory target site, and subsequently examined whether the resulting circuit could modulate CTNNB1 and downstream Wnt-associated gene expression during differentiation.

## 2. Results

### 2.1 Transient TBXT induction identifies an early differentiation window for promoter linked regulation

We designed a regulatory configuration in which a developmental promoter drives an intracellular effector during a defined differentiation stage (Figure 1A). The T/Brachyury promoter was selected to connect effector expression to an early mesendoderm-associated state. To establish the corresponding expression window in H9 cells, differentiation was initiated with a CHIR99021 pulse and endogenous TBXT expression was followed over the first five days. TBXT increased sharply at the beginning of differentiation, peaked on day 1, remained elevated on day 2, and declined thereafter (Figure 1B). Because this expression window coincided with the initial lineage transition, we reasoned that the corresponding promoter sequence could serve as a stage-responsive temporal input to restrict downstream effector expression to early differentiation.

**Figure 1.**
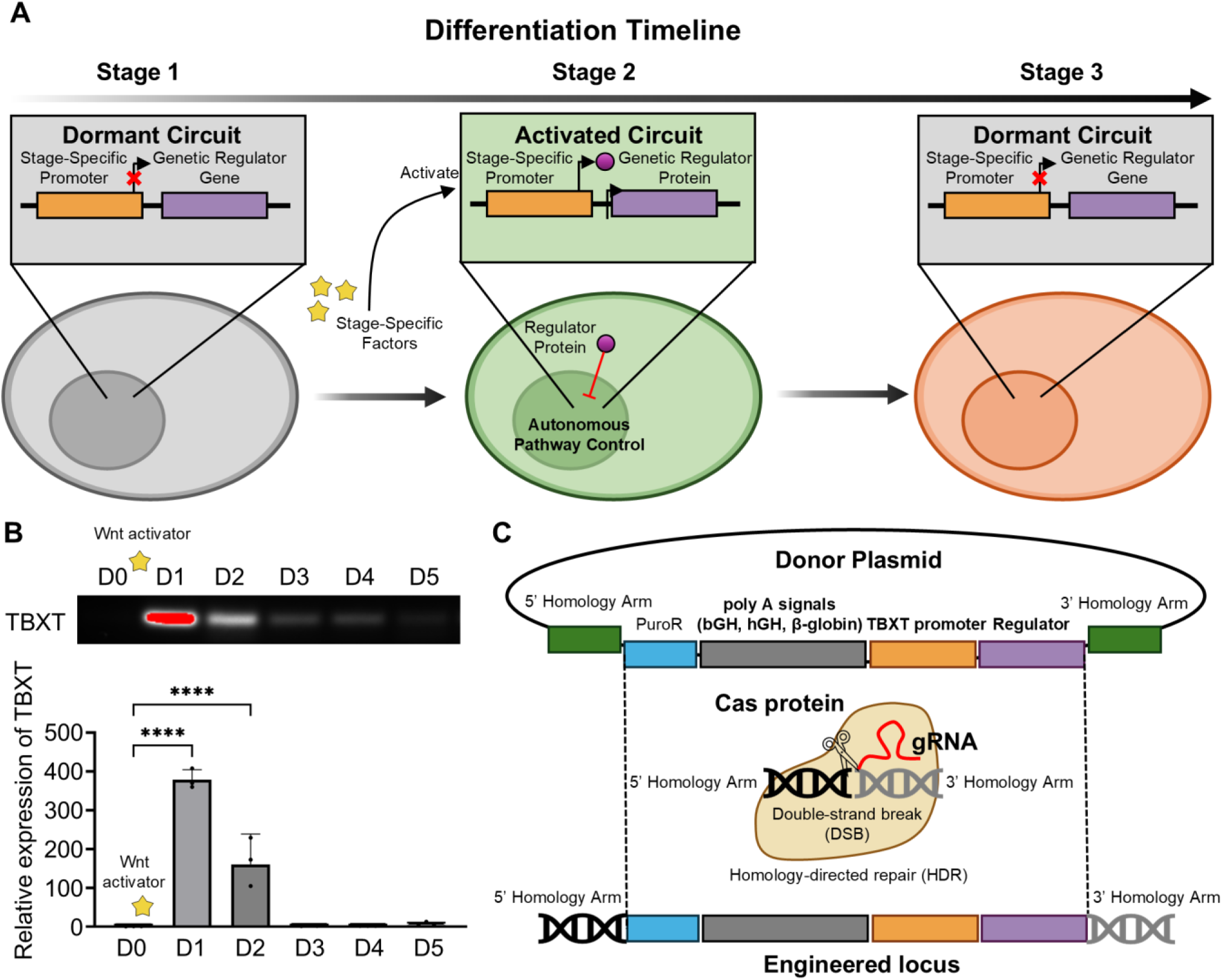
Developmental promoter strategy and early TBXT expression. (A) Conceptual design of a developmental promoter driving a genetic regulator during a defined differentiation stage. The illustrated dormant and activated states depict the intended temporal control of regulator expression. (B) Endogenous TBXT expression in H9 human pluripotent stem cells during days 0–5 of differentiation, assessed by endpoint RT-PCR (top) and quantitative PCR (bottom). Cells received 12 μM CHIR99021 from D0 to D1; D0 samples were collected before treatment. Quantitative expression was normalized to RPL13A and scaled to the mean D0 level. Bars show mean ± SD, with individual points representing three separately differentiated wells (n = 3). One-way ANOVA followed by Tukey multiple comparisons; ****P < 0.0001 for the indicated comparisons. (C) AAVS1-targeting donor architecture and homology-directed integration strategy. The donor contains a promoterless puromycin resistance cassette, tandem bovine growth hormone, human growth hormone, and β-globin polyadenylation signals upstream of the T promoter, and a downstream reporter or regulatory gene flanked by homology arms. The polyadenylation elements were included to limit upstream transcriptional read-through. The promoter fragment used in the donor is murine T/Brachyury; endogenous human Brachyury expression is designated TBXT.

To implement this strategy, we designed a donor for integration at the AAVS1 site within PPP1R12C, containing a promoterless puromycin selection cassette followed by the T-promoter-driven reporter or regulator (Figure 1C). The selection cassette was designed to obtain puromycin resistance through transcription driven by the endogenous PPP1R12C promoter. To limit potential read-through into the downstream circuit, three tandem polyadenylation signals were placed between the selection cassette and the T promoter. This arrangement was intended to limit host-derived transcription while providing a shared framework for evaluating the promoter with EGFP and subsequently coupling it to a gene-regulatory effector.

### 2.2 The integrated T promoter drives a transient reporter response during differentiation

Having established the endogenous TBXT expression window, we next asked whether the integrated T-promoter fragment would show a corresponding differentiation-associated response. We therefore generated a T-EGFP reporter line (Figure 2A). For downstream analysis, an isolated T-EGFP clone was selected and verified by junction PCR, which confirmed the expected 5′ and 3′ integration products alongside the absence of an amplicon from the unmodified locus (Supplementary Figure 1). Following differentiation induction, EGFP was detected by immunoblotting and fluorescence microscopy (Figure 2B,C). The immunoblot showed the clearest EGFP signal on day 2.

**Figure 2.**
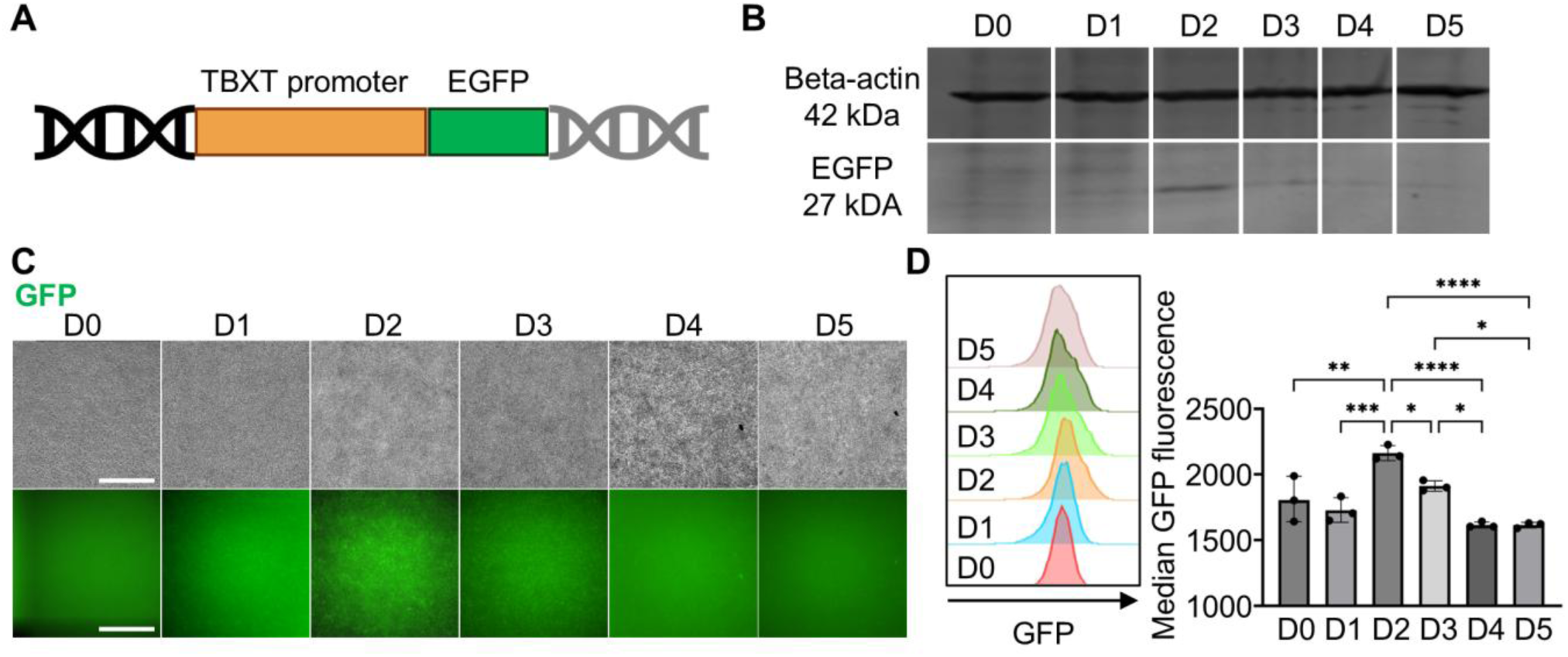
Transient T promoter reporter activity during differentiation. (A) Schematic of the T-promoter-driven EGFP reporter integrated at AAVS1. (B) Immunoblots for β-actin (42 kDa) and EGFP (27 kDa) in T-EGFP cells during days 0–5 of differentiation; β-actin served as the loading control. (C) Brightfield images (top) and corresponding GFP fluorescence images (bottom) of T-EGFP cultures at the indicated time points. Scale bars = 500 µm. (D) Representative live-cell GFP fluorescence distributions (left) and quantification of GFP fluorescence (right). Each point represents the median fluorescence within the GFP gate for one separately differentiated well. Cells received CHIR99021 from D0 to D1, and D0 samples collected before CHIR treatment served as the pre-induction control. Bars show mean ± SD, n = 3 wells per time point. One-way ANOVA followed by Tukey multiple comparisons; *P < 0.05, **P < 0.01, ***P < 0.001, and ****P < 0.0001 for the indicated comparisons.

Live-cell flow cytometry similarly identified a transient increase in reporter fluorescence, with the highest measured GFP intensity on day 2 and lower intensity at later time points, relative to pre-induction (day 0) controls (Figure 2D). Notably, the timing of this reporter peak lagged slightly behind the initial surge of endogenous TBXT, consistent with promoter activation followed by subsequent reporter protein accumulation. These complementary measurements support the utility of the integrated T-promoter fragment as a developmentally responsive expression element in this differentiation setting.

### 2.3 Cas12a mediated integration avoids detectable CTNNB1 target site indels

Having verified the reporter response, we next replaced EGFP with dCas9-KRAB alongside a U6-driven guide cassette targeting CTNNB1 to create the T-dCas9-CTNNB1 construct (Figure 3A). In this configuration, the timing of repression is governed by the developmental promoter, which drives dCas9-KRAB only during early differentiation, whereas the guide RNA is constitutively expressed under the U6 promoter to direct the repressor specifically to the CTNNB1 locus.

**Figure 3.**
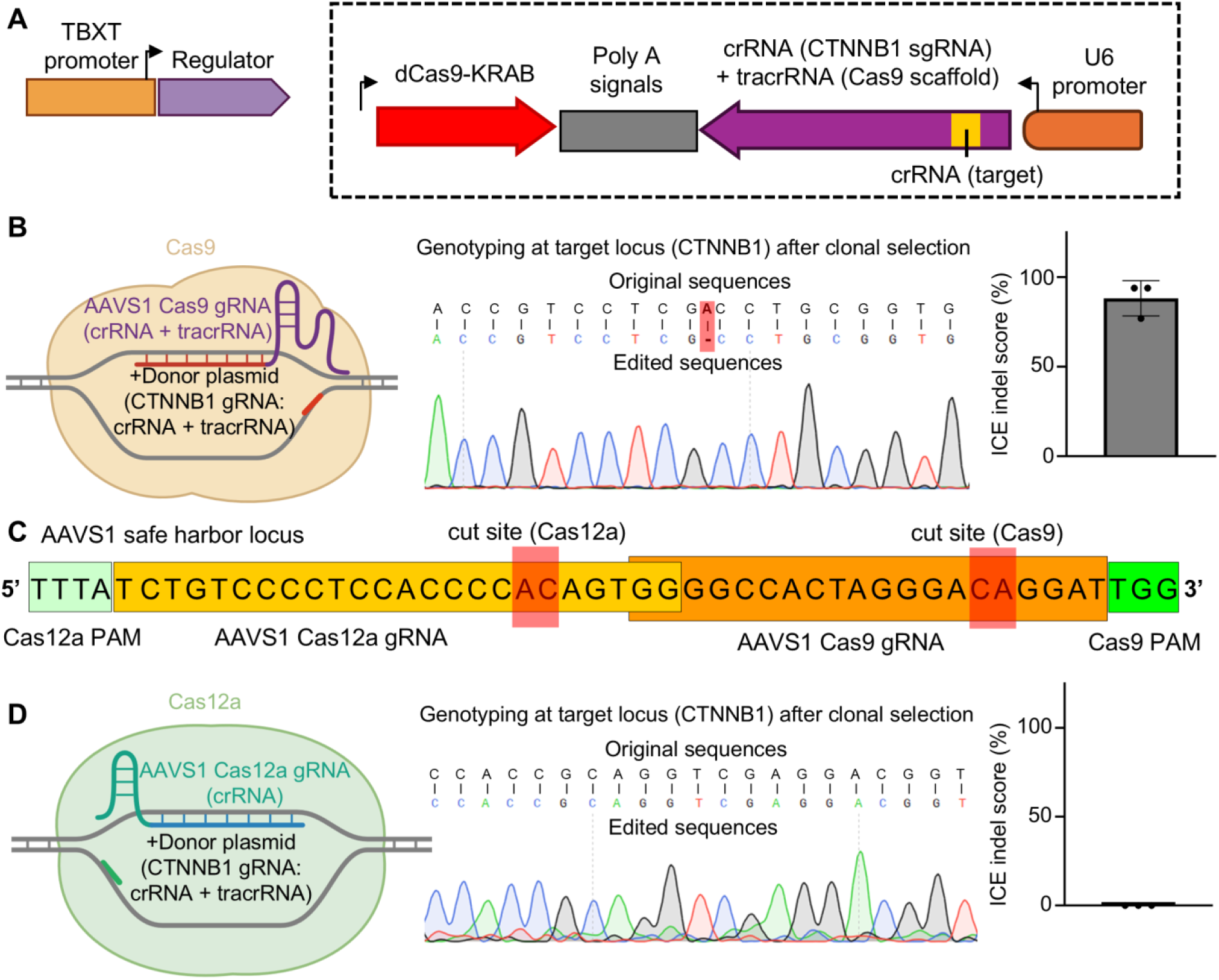
Cas12a mediated integration avoids detectable CTNNB1 target site indels. (A) Regulatory cassette containing T-promoter-driven dCas9-KRAB and a U6-driven CTNNB1-targeting single-guide RNA, comprising a target-specific spacer and the Cas9 guide scaffold. The U6 cassette is oriented opposite to T-promoter-driven transcription. (B) Cas9-based AAVS1 integration workflow (left), representative Sanger sequencing at the CTNNB1 guide target region after clonal selection (middle), and ICE-estimated indel percentages (right). The highlighted alignment indicates a deletion in the representative Cas9-derived clone. (C) AAVS1 sequences targeted by the Cas9 and Cas12a integration workflows, showing their guide-binding regions, protospacer-adjacent motifs (PAMs), and indicated cleavage sites. (D) Cas12a-based AAVS1 integration workflow (left), representative CTNNB1 target-region sequencing after clonal selection (middle), and ICE-estimated indel percentages (right). ICE scores were obtained from the original analysis of edited-sample Sanger traces relative to an unmodified H9 control. Each point represents a distinct clone, and bars show mean ± SD for three clones per workflow (n = 3). The three Cas9-derived clones were randomly selected. All three examined Cas12a-derived clones had ICE scores of zero, indicating no indel contribution detected at the assayed CTNNB1 target site by this analysis.

However, this assembly introduced a potential technical complication because the active Cas9 nuclease supplied for AAVS1 integration could inadvertently complex with the donor-encoded CTNNB1 guide RNA and cleave the target locus. Consistent with this concern, clones generated using the Cas9 workflow showed unintended indels at the CTNNB1 target site, with Sanger sequencing and ICE analysis confirming substantial indel frequencies across all three randomly screened clones (Figure 3B). ICE estimates the percentage of sequences containing indels by decomposing an edited Sanger trace relative to a control trace [19,20]. Such permanent sequence changes at a site intended solely for transcriptional repression underscored the need for an alternative integration strategy.

To circumvent this cross-reactivity, we switched to a Cas12a ribonucleoprotein workflow to direct AAVS1 integration at a nearby target site (Figure 3C, D). Cas12a employs a distinct guide architecture from Cas9 [21], providing a means to integrate the donor without reusing the nuclease associated with its regulatory guide. The three examined Cas12a-derived clones exhibited ICE scores of zero at the CTNNB1 target site, in contrast to the Cas9-derived clones (Figure 3D). Thus, the Cas12a-based workflow yielded engineered clones without detectable CTNNB1 target-site indels in our analysis. Furthermore, junction PCR performed on the representative clone selected for downstream differentiation assays confirmed targeted integration, showing the expected 5′ and 3′ products alongside no detectable amplification from the unmodified locus (Supplementary Figure 1).

### 2.4 The T promoter CRISPRi line shows CTNNB1 repression and attenuated Wnt associated transcription

Following Cas12a-mediated integration, we evaluated whether the T-dCas9-CTNNB1 circuit modulated target gene expression during differentiation. When subjected to directed differentiation in parallel with parental H9 cells, the engineered line exhibited a marked reduction in CTNNB1 transcript levels, reaching statistically significant downregulation on days 3 and 4 (Figure 4A). To determine whether this targeted repression altered downstream Wnt pathway activity, we analyzed four canonical target genes: SP5, TCF7, LEF1, and AXIN2. All four downstream targets showed blunted induction profiles relative to controls, though with distinct kinetics (Figure 4B). While SP5 displayed an immediate attenuation during early induction (day 1), TCF7, LEF1, and AXIN2 maintained lower expression across subsequent differentiation stages. Together, the down-regulation of CTNNB1 and the corresponding suppression of downstream Wnt-responsive transcripts confirm that the integrated circuit effectively attenuates pathway activation during early cell-fate specification

**Figure 4.**
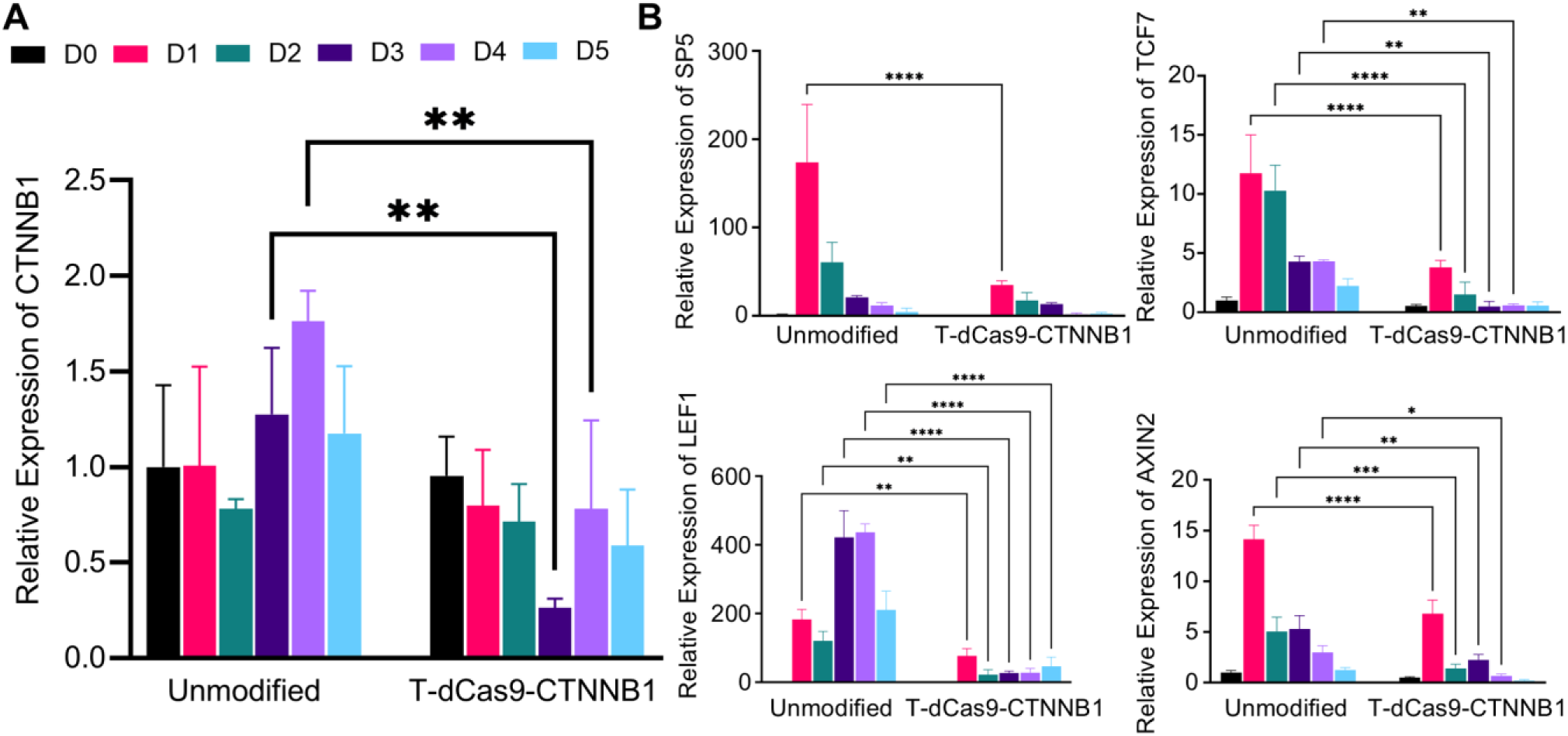
CTNNB1 repression and attenuation of Wnt associated transcription. Quantitative PCR analysis of (A) CTNNB1 and (B) SP5, TCF7, LEF1, and AXIN2 expression in unmodified H9 cells and the T-dCas9-CTNNB1 line during days 0–5 of differentiation. Both lines were cultured under matched conditions, with 12 μM CHIR99021 treatment from D0 to D1 in RPMI 1640 supplemented with B27 without insulin. D0 samples were collected before CHIR treatment. Expression was normalized to RPL13A and scaled to the mean unmodified H9 D0 level. Data are presented as mean ± SD from three biological replicates (n = 3). Each gene was analyzed separately by two-way ANOVA with cell line, differentiation day, and their interaction, followed by Šidák-adjusted comparisons between the two cell lines at each time point. The six time-point comparisons formed one multiple-comparison family per gene. *P < 0.05, **P < 0.01, ***P < 0.001, and ****P < 0.0001 for the indicated comparisons.

## 3. Discussion

This study explores a modular strategy for connecting intracellular gene regulation to developmental state. The results establish three elements of that strategy: a T-promoter reporter with a transient differentiation-associated response, an integration workflow that avoids detectable indels at the intended CRISPRi target, and reduced CTNNB1 expression accompanied by attenuation of Wnt-associated transcription in the selected engineered line. Overall, these findings provide an initial proof of concept that an endogenous developmental promoter can be coupled to a programmable intracellular regulator to influence signaling pathways during hPSC differentiation.

The motivation for this approach is to connect the timing of an intervention to the state of the responding cell. Extracellular factors remain powerful tools for initiating and directing differentiation, but their effects depend on cellular context and the endogenous signaling environment [1,4]. A promoter responsive to an emerging developmental state could provide an additional layer of coordination: cells that enter the relevant state would begin expressing an encoded regulator. In the present system, CHIR initiates differentiation, while the T promoter provides the proposed link between the resulting transcriptional state and effector expression. This design complements externally controlled differentiation and creates a route toward interventions that follow developmental progression within the cell.

The T promoter is particularly informative because its activity is associated with an early developmental transition and is connected to Wnt/β-catenin signaling [17,18]. Consistent with this role, our transient reporter response confirms that the integrated promoter fragment faithfully retains its differentiation-associated activity. Coupling this responsive input to a CTNNB1-directed repressor successfully creates an autonomous negative-feedback loop, as evidenced by the concurrent reduction of CTNNB1 transcripts and attenuation of Wnt-responsive gene induction. Importantly, while the circuit effectively dampened overall pathway activation, the onset and magnitude of repression varied across individual downstream targets, pointing to complex attenuation kinetics following initial promoter activation. Notably, downstream Wnt targets such as SP5 and AXIN2 displayed attenuated induction as early as day 1, preceding the statistically significant downregulation of CTNNB1 observed on days 3 and 4. This temporal offset likely reflects differences in transcript turnover and locus-specific responsiveness: while CTNNB1 is an abundant, constitutively transcribed component critical for cellular homeostasis whose pre-existing mRNA pool takes time to substantially deplete [22], downstream targets such as SP5 and AXIN2 are immediate early-response genes whose de novo induction is acutely sensitive to even subtle reductions in initial Wnt signaling flux [23,24]. Early dCas9-KRAB accumulation may thus dampen nascent CTNNB1 transcription sufficiently to blunt downstream activation thresholds before total CTNNB1 steady-state levels decline to statistical significance

Beyond dynamic pathway control, another practical contribution of this study is decoupling circuit integration from circuit function. While the donor cassette was designed exclusively for transcriptional repression, its encoded U6-CTNNB1 guide RNA can inadvertently pair with the active Cas9 nuclease supplied for genomic integration, causing unintended target-site indels as observed in our initial workflow. To overcome this cross-reactivity, we adopted a Cas12a-based integration strategy, which successfully yielded correctly integrated clones without detectable indels at the target locus. These findings demonstrate the necessity of employing mutually orthogonal nuclease–guide systems when delivery and regulatory machineries risk overlapping. This general design principle should prove valuable for engineering other stable, multi-component CRISPR circuits without unintended genomic editing.

A central design consideration for cell-state-linked circuits is ensuring that the transcriptional strength of the developmental promoter matches the operational threshold of the chosen effector. Although the T promoter drove relatively low to modest reporter levels, this output was sufficient to mediate measurable functional repression, highlighting that sustained or high-level expression is not strictly required for effective CRISPR interference. Indeed, even promoters with modest transcriptional output can provide effective regulation when paired with potent transcriptional repressors [25] or epigenome modifiers that establish self-sustaining transcriptional memory [11]. Circuit responsiveness could be further calibrated by systematically tuning effector turnover rates and promoter sensitivities across different differentiation thresholds. Building upon these tunable single-node circuits, this approach could be expanded to orchestrate complex differentiation trajectories. Because synthetic lineage-control networks and direct transcription factor programming have already proven capable of redirecting developmental fates [12, 26], coupling multiple stage-specific promoters to distinct regulatory effectors could enable sequential or combinatorial control across successive lineage transitions.

While these findings establish proof of concept, several technical aspects warrant further refinement. Because functional characterization was conducted in a representative clonal line, analyzing additional independently isolated clones alongside non-targeting guide controls and target rescue will be critical to rigorously confirm guide specificity. In addition, directly profiling dCas9-KRAB and β-catenin protein kinetics, paired with live Wnt signaling reporters, will help resolve the quantitative lag between promoter activation and downstream pathway inhibition. It is also worth noting that our comparison between integration strategies encompassed both nuclease type (Cas9 vs. Cas12a) and delivery format (plasmid vs. RNP); isolating these variables would provide deeper mechanistic insight into nuclease cross-reactivity during donor delivery. Ultimately, assessing whether this autonomous regulatory strategy improves terminal lineage yield or buffers against batch-to-batch variation will be crucial for evaluating its utility in scalable cell manufacturing.

In conclusion, the present findings establish a practical starting point for that development: a validated developmental promoter input, a compatible integration strategy, and a measurable transcriptional response from the linked regulatory construct. Expanding the repertoire of promoter–effector combinations could make cell-state-responsive regulation a useful component of hPSC differentiation engineering.

## 4. Materials and methods

### 4.1 Cell culture and differentiation

H9 human embryonic stem cells (WiCell) were maintained on Matrigel-coated cell culture plates in mTeSR1 (STEMCELL Technologies, 85850) at 37 °C and 5% CO₂. Cells were routinely passaged with Versene (Gibco, 15040066) and dissociated with Accutase (STEMCELL Technologies, 07922) for differentiation experiments. Differentiation was initiated on day 0 at 70–80% cell confluency by treatment with 12 μM CHIR99021 (CHIR / Selleck Chemicals, S1263) in RPMI 1640 medium (Gibco, 11875093) supplemented with insulin-free B27 (B27- / Gibco, A1895601). CHIR was removed on day 1. RPMI 1640 with B27- was supplied on days 0, 1, 3, and 5. Engineered and unmodified cells used for the Figure 4 comparison were cultured under matched conditions. Day 0 samples were collected before CHIR treatment and served as the pre-induction control.

### 4.2 Donor construction

Donor construction began with AAVS1-Pur-CAG-HESX1, a derivative of AAVS1-Pur-CAG-mCherry (Addgene #80946). The starting plasmid was linearized with SpeI-HF (NEB, R3133S) and MluI-HF (NEB, R3198S) to obtain the AAVS1 donor backbone. A murine T/Brachyury promoter fragment, based on the previously reported reporter sequence [18], was synthesized as a gBlock (Integrated DNA Technologies). Additional polyadenylation elements were amplified from the starting donor and assembled with the promoter-containing insert using In-Fusion HD cloning (Takara Bio, 102518). The resulting donor architecture contained a puromycin resistance cassette followed by three tandem polyadenylation signals (bovine growth hormone, human growth hormone, and β-globin poly(A)) placed upstream of the T promoter to insulate against upstream transcriptional read-through. A promoter-containing intermediate was remodeled using PspXI (NEB, R0656S) and EcoRV-HF (NEB, R3195S) to provide a cloning site downstream of the T promoter. For the CRISPRi donor, the backbone was linearized with EcoRV-HF and assembled with dCas9-KRAB amplified from pLV hU6-sgRNA hUbC-dCas9-KRAB-T2a-Puro (Addgene #71236) and a synthesized U6–guide RNA cassette. The dCas9-KRAB insert included an N-terminal 3×FLAG tag, nuclear localization sequences, and a terminal stop codon introduced by the reverse amplification primer; WPRE was retained downstream of the coding region. In the final CRISPRi donor, the U6–guide cassette was oriented opposite to T-promoter-driven transcription. The CTNNB1 gRNA spacer was 5′-GTCCGACCGTCCTCGACCTG-3′, excluding the adjacent genomic PAM. The T-EGFP donor placed a Kozak sequence and EGFP downstream of the same T-promoter input. Both donor constructs contained 804-bp left and 837-bp right homology arms designed for homology-directed integration at AAVS1, a locus previously validated for targeted transgene insertion in hPSCs [22].

### 4.3 Generation of engineered cell lines

H9 cells were pretreated with 10 µM Y-27632 (Selleck Chemicals, S1049) overnight before dissociation. For electroporation, approximately 5 × 10⁶ cells were harvested and prepared in a total volume of 400 µL PBS. For the Cas9 workflow, 6 μg Cas9-GFP plasmid (Addgene #44719), 4 μg AAVS1 gRNA T2 plasmid (Addgene #41818), and 10 μg donor plasmid were combined with cells in PBS. For the Cas12a workflow, ribonucleoprotein (RNP) complexes were assembled by incubating 1.6 nmol AAVS1-targeting crRNA and 0.315 nmol Cas12a protein (Integrated DNA Technologies) at room temperature for 10–20 min. The assembled RNP was combined with cells, 10 μg donor plasmid, and 10 μL of 78 μM electroporation enhancer (Integrated DNA Technologies, 1076300). The Cas12a target site was located adjacent to the Cas9 target at the AAVS1 locus (Figure 3C). The cell suspension was divided equally between two 0.4-cm cuvettes (Bio-Rad, 1652091) and electroporated using an electroporator (Bio-Rad, Gene Pulser Xcell) at 320 V, 200 µF, and 1,000 Ω. Following electroporation, cells were seeded onto Matrigel-coated plates in mTeSR1 supplemented with 10 µM Y-27632. After at least one week of recovery, stably integrated cells were selected with 1 µg/mL puromycin. Individual drug-resistant colonies were manually picked and expanded for genotyping. The verified CTNNB1-targeting clone used for downstream functional differentiation experiments was designated T-dCas9-CTNNB1.

### 4.4 Genotyping and ICE analysis

Genomic DNA was isolated with the Quick-DNA Miniprep Plus kit (Zymo Research, D4068). PCR was performed using GoTaq Green Master Mix (Promega, M712). Primer pairs interrogating the 5′ and 3′ insertion junctions and the unmodified AAVS1 locus were used to assess integration (Supplementary Figure 1; Supplementary Table 1). CTNNB1 target-region amplicons were analyzed by Sanger sequencing at Functional Biosciences. Indel estimates were obtained using the web-based Inference of CRISPR Edits service (ICE; currently provided by EditCo Bio, Inc., https://www.editco.bio) by comparing edited-sample and unmodified-control Sanger traces [19,20]. The ICE score reports the estimated percentage of sequences containing insertions or deletions, including edits regardless of their predicted functional consequence, as defined in Understanding your ICE Export. An ICE score of zero denotes no indel contribution detected by the analysis at the assayed site. Figure 3 reports the original service-generated ICE scores for three randomly selected Cas9-derived clones and three Cas12a-derived clones, labeled C1–C3 within each group.

### 4.5 RNA analysis

RNA was extracted using TRIzol reagent (Ambion, 15596026) and the Direct-zol RNA Miniprep kit (Zymo Research, R2050), and cDNA was prepared using the Omniscript reverse transcription kit (Qiagen, 205111). Endpoint RT-PCR used GoTaq Green Master Mix. Quantitative PCR was performed with PowerUp SYBR Green Master Mix (Applied Biosystems, 4367659) on an AriaMx Real-Time PCR System (Agilent, G8830A). Target Cq values were normalized to the RPL13A reference assay. Relative expression was calculated from reference-normalized expression and scaled to the mean of the unmodified H9 D0 samples. TBXT was measured to characterize the early differentiation response; CTNNB1, SP5, TCF7, LEF1, and AXIN2 were measured to assess the engineered line. Sequences and expected amplicon sizes for the RPL13A, TBXT, CTNNB1, SP5, TCF7, LEF1, and AXIN2 assays are provided in Supplementary Table 2.

### 4.6 Reporter imaging and flow cytometry

Live T-EGFP cultures were imaged using a Nikon Eclipse Ti2-E fluorescence microscope with a temperature-controlled stage (Tokai Hit, Tokai Hit STX Series Stage Top Incubator System). For flow cytometry, cells were detached at the indicated time points and analyzed as live cells on a BD Accuri C6 Plus. GFP fluorescence was measured directly and analyzed in FlowJo.

### 4.7 Immunoblotting

Cells were lysed in 1× RIPA buffer (Rockland, MB-030-0050) supplemented with 1× protease inhibitor cocktail (Thermo Scientific, 78430), incubated at 4 °C for 5 min, and clarified by centrifugation at 15,000 × g for 5 min at 4 °C. Total protein concentrations were determined using a BCA protein assay kit (Thermo Scientific, 23225). Equal amounts of protein were separated on Novex WedgeWell 4–12% Tris-Glycine gels (Invitrogen, XP04125BOX) under denaturing conditions and transferred onto nitrocellulose membranes. Membranes were blocked with 5% non-fat dry milk (Santa Cruz Biotechnology, sc-2324) in TBST for 1 h at room temperature and subsequently incubated overnight at 4 °C with shaking with the following primary antibodies diluted in blocking buffer: mouse anti-GFP (Invitrogen, MA5-15256; 1:1,000) and rabbit anti-β-actin (Cell Signaling Technology, 4967S; 1:1,000). After washing, membranes were incubated for 1 h at room temperature in the dark with IRDye-conjugated secondary antibodies diluted in blocking buffer: goat anti-mouse IRDye 680RD (LI-COR, 926-68070) and goat anti-rabbit IRDye 800CW (LI-COR, 926-32211). Blots were washed and imaged on an Odyssey Imaging System (LI-COR).

### 4.8 Statistical analysis

Data are presented as mean ± standard deviation. For Figures 1B, 2D, and 4, n = 3 represents three biological replicates per condition and time point. For Figure 3, n = 3 represents three distinct clones per integration workflow. Figures 1B and 2D were analyzed using ordinary one-way ANOVA followed by Tukey multiple comparisons. Figure 4 was analyzed separately for each gene using ordinary two-way ANOVA with cell line, differentiation day, and their interaction, followed by Šidák-adjusted comparisons of the two cell lines at each time point. The six time-point comparisons constituted one multiple-comparison family per gene. Statistical significance was defined as P < 0.05; symbols denote *P < 0.05, **P < 0.01, ***P < 0.001, and ****P < 0.0001.

## 5. Acknowledgments

This work was supported by the National Science Foundation (NSF) through the Engineering Research Center for Cell Manufacturing Technologies (CMaT; EEC-1648035) and the NSF RECODE program (Award No. 2225300).

