## Supplementary information for "Coupling a developmental promoter to CRISPR interference for Wnt pathway regulation in human pluripotent stem cells"

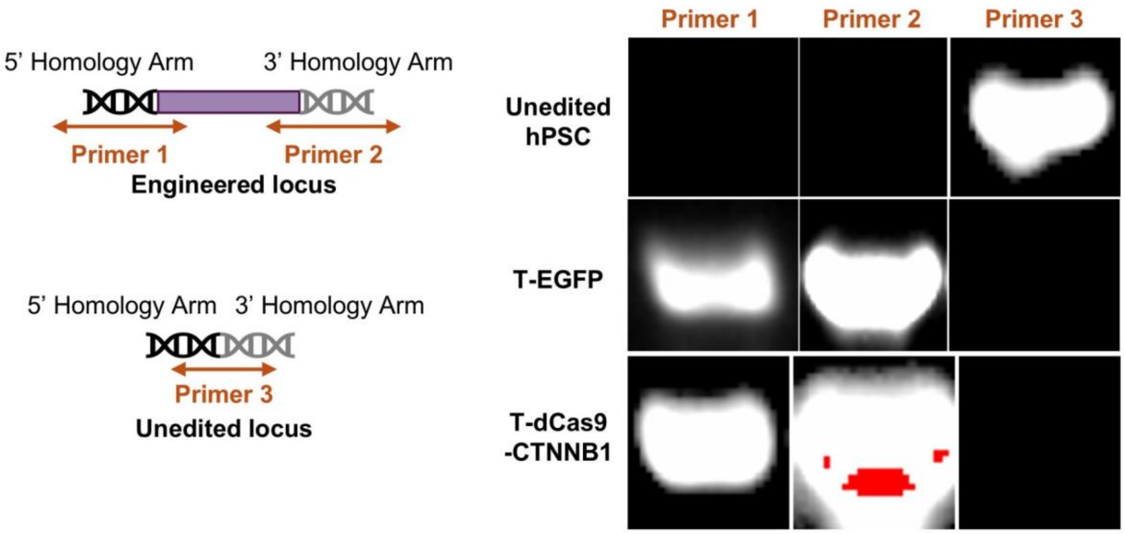


**Supplementary Figure 1. PCR assessment of targeted integration**. Schematic primer-pair positions and PCR products at the engineered and unmodified AAVS1 loci. Primer pairs 1 and 2 interrogate the 5′ and 3′ insertion junctions, respectively; primer pair 3 interrogates the unmodified locus. Following puromycin selection, colonies were picked and screened by PCR. A representative clone from each engineered line (T-EGFP and T-dCas9-CTNNB1) is shown alongside parental H9 cells; the engineered clones yielded the 5′ and 3′ integration products, with no detectable product from the unmodified-locus assay. Primer sequences are provided in Supplementary Table 1.

**Supplementary Table 1. Primers for AAVS1 integration screening**

| Primer pair and assay | Forward sequence 5′ to 3′ | Reverse sequence 5′ to 3′ | Expected size bp |
|---|---|---|---|
| 1 5′ junction | GAACTCTGCCCTCTAACGCT | CGTCACCGCATGTTAGAAGA | 1002 |
| 2 3′ junction | TAAAGCCTGGGGTGCCTAAT | CTTCTTGGCCACGTAACCTG | 1220 |
| 3 Unmodified locus | TGCTTTCTTTGCCTGGACAC | TCTGGGCGGAGGAATATGTC | 1000 |

Expected sizes for primer pairs 1 and 2 refer to insertion-associated junction products; the size for primer pair 3 refers to the unmodified allele.

**Supplementary Table 2. Primers for gene expression analysis**

| Gene | Forward sequence 5′ to 3′ | Reverse sequence 5′ to 3′ | Expected size bp |
|---|---|---|---|
| TBXT | GAACGGCAGGAGGATGTTTC | AGGAAGGAGTACATGGCGTT | 75 |
| RPL13A | CGTGCGTCTGAAGCCTACAA | CCGTAGCCTCATGAGCTGTTTC | 157 |
| SP5 | GAGTTCTCGCCGGTCAAGAT | GAGGCAGCAGGTTGGAGTAG | 124 |
| TCF7 | GTGCACACTTAAGGAGAGCG | CTTGGTGCTTTTCCCTCGAC | 194 |
| LEF1 | ACGAGCACTTTTCTCCAGGA | CAAGAGGTGGGGTGATCTGT | 150 |
| AXIN2 | GGTCCACGGAAACTGTTGAC | TCCATCTACACTGCTGTCCG | 179 |
| CTNNB1 | AGCTGGCCTGGTTTGATACT | TCAGCAACTCTACAGGCCAA | 87 |
